## Supplementary material for "The Jag2/Notch1 signaling axis promotes sebaceous gland differentiation and controls progenitor proliferation": All supplemental figures revised

**Figure S1. Sebaceous ducts remain unaffected after Notch inhibition.**

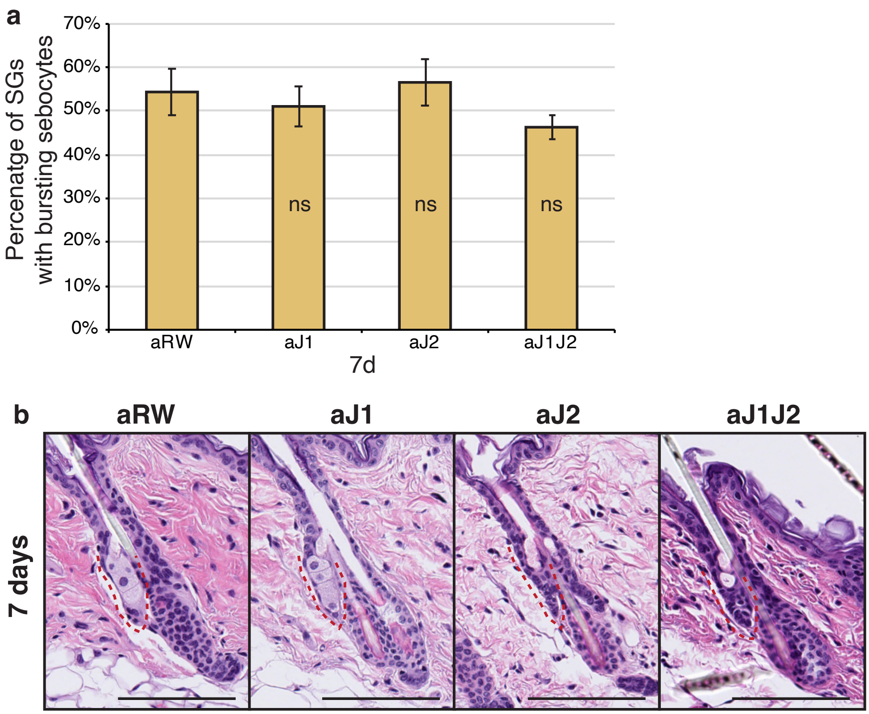

**Figure S2. Notch is active in the sebaceous gland stem cells.**

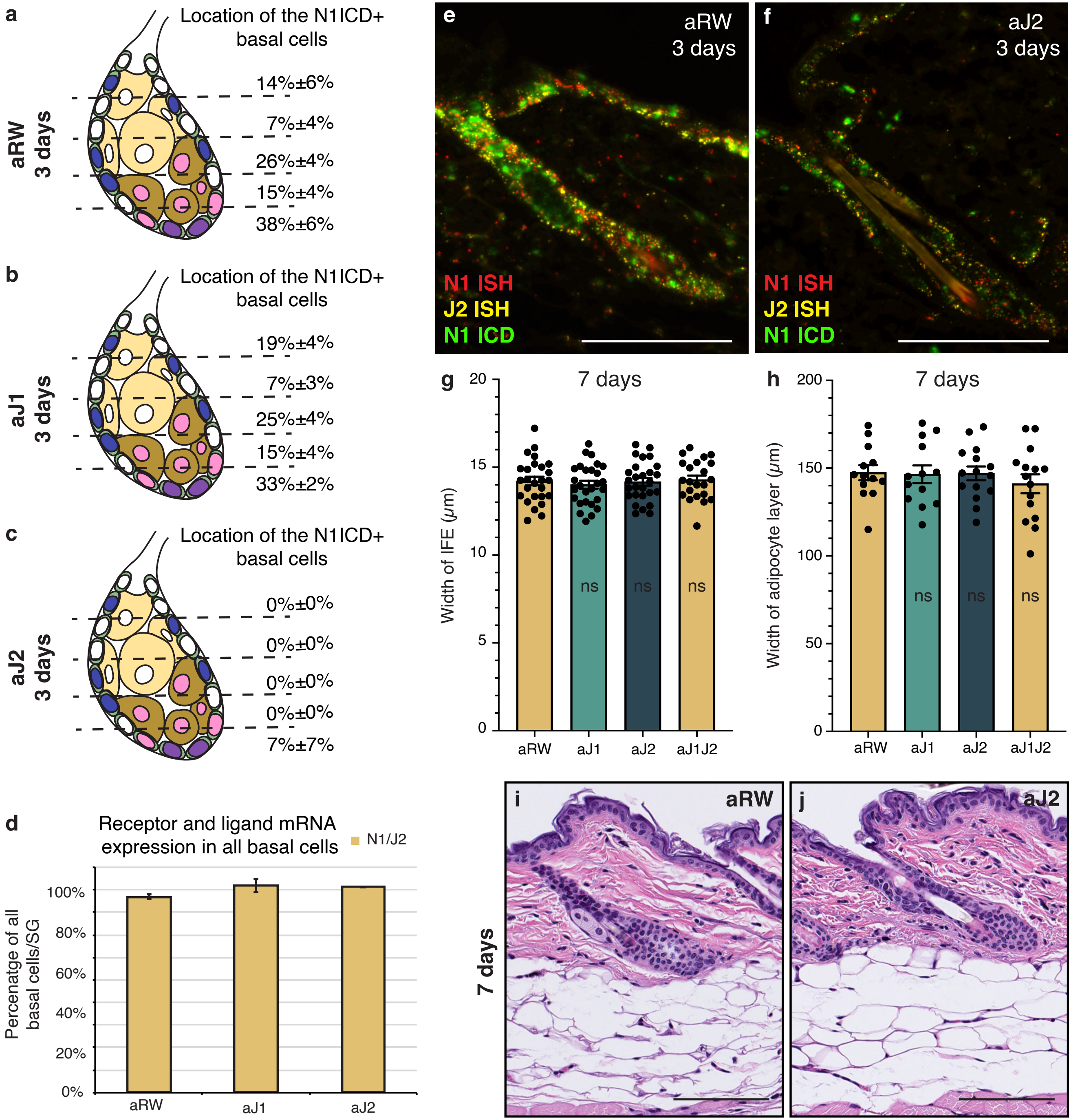

**Figure S3. Loss of Notch activity in the SG stem cells inhibits sebocyte differentiation.**

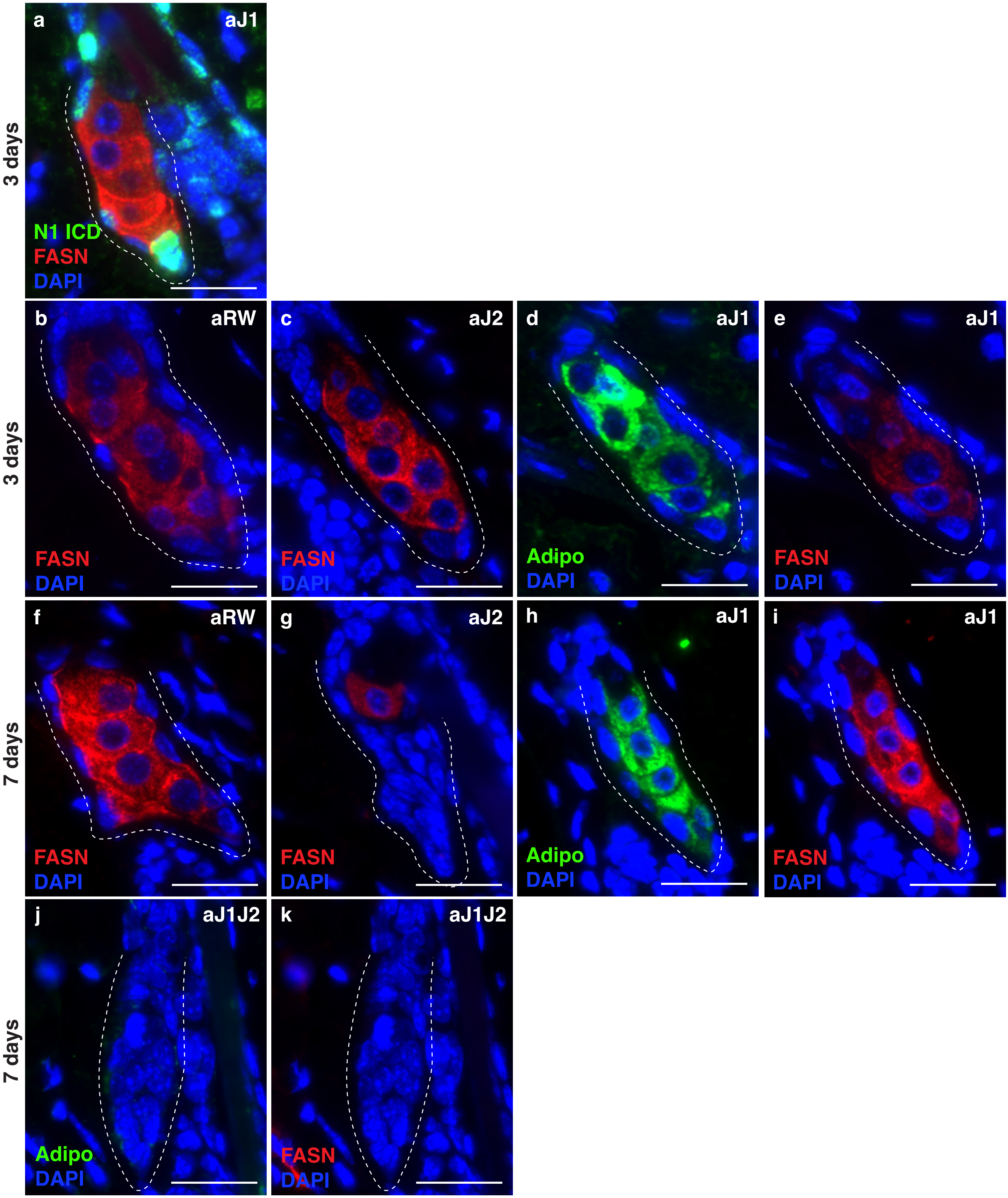

**Figure S4. Notch activity in the SG stem cells is required to prevent unregulated progenitor proliferation.**

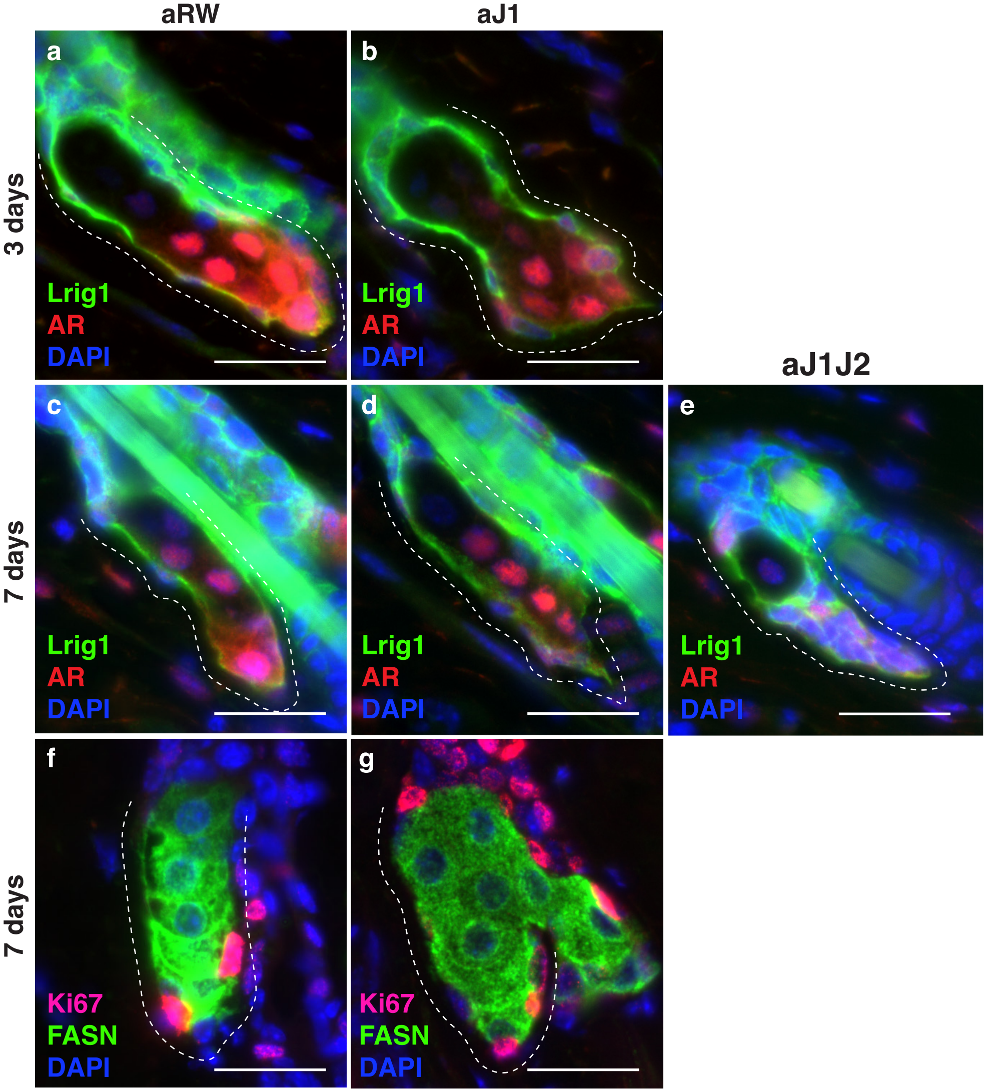

**Figure S5. The block in sebocyte differentiation is lifted upon recovery of Notch activity.**

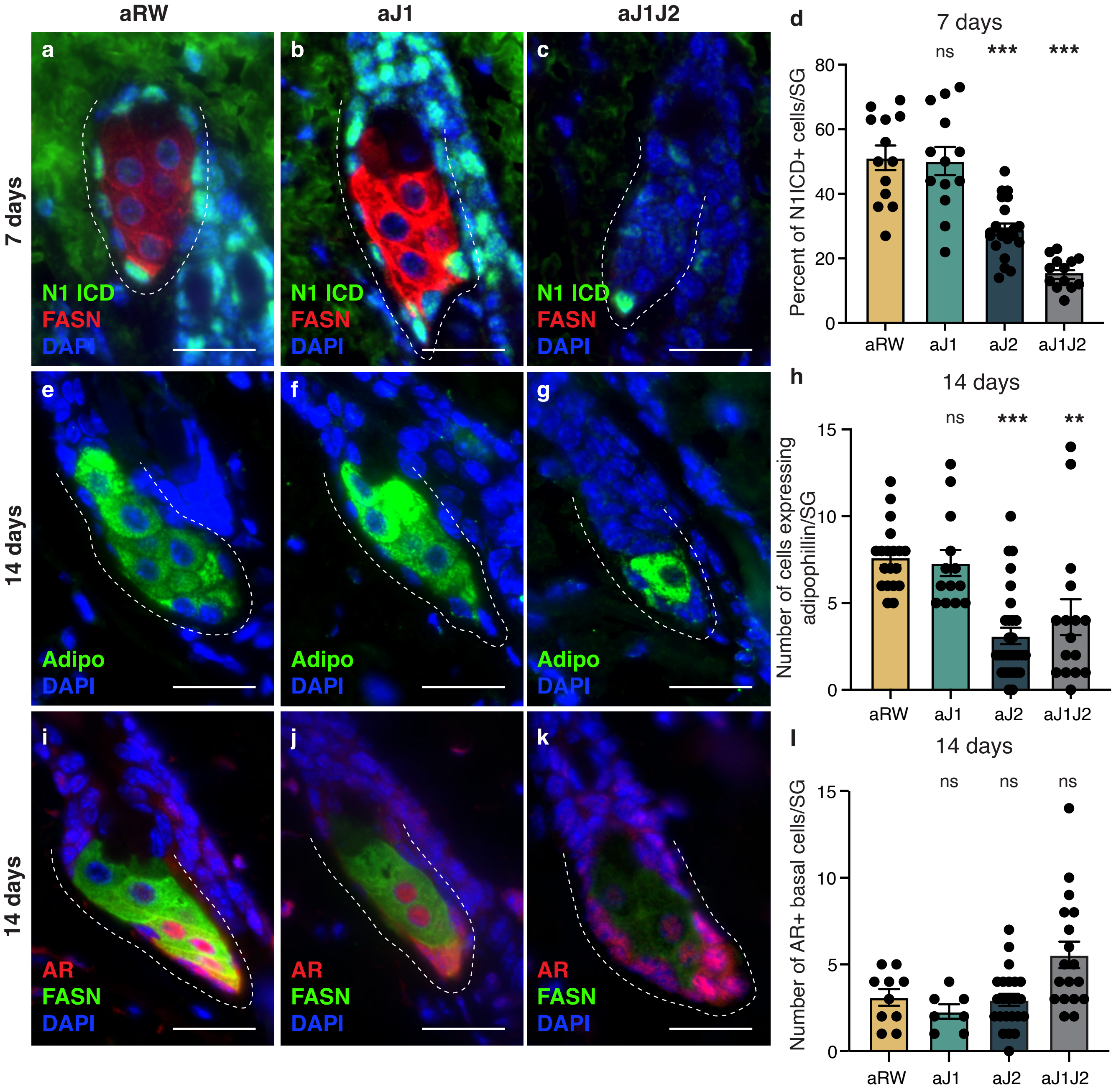
